# Smooth move: Behavioural changes in captive western lowland gorillas (*Gorilla gorilla gorilla*) after group split and relocation

**DOI:** 10.64898/2026.06.25.734437

**Authors:** Marie Žabková, Monika Čížková, Vedrana Šlipogor, Roman Vodička, Martina Konečná

## Abstract

Zoological gardens strive to prioritize excellent animal care and adhere to the highest standards of animal welfare, ensuring that the conditions of animals’ social and physical environment are as close as possible to those in the wild. This study investigates the effects of group split and relocation to the new enclosure on the behaviour of western lowland gorillas (*Gorilla gorilla gorilla*) (N = 7) living in Prague Zoo, Czech Republic. We conducted over 289 hours of behavioural observations focusing on daily activities, social interactions, and behaviours that serve as potential stress and welfare indicators. The group split led to establishment of two groups in two separate enclosures; the old enclosure consisted of a ‘bachelor group’ (i.e. three males) and a new enclosure consisted of a mixed-sex group (i.e. three females and one juvenile male). The behavioural comparisons across different study periods were conducted using linear mixed models (LMMs). The changes led to an increase in time spent moving, feeding, being in social proximity, and higher rates of approaches among the gorillas, as well as to a decrease in rates of self-directed and ‘undesirabl’ behaviors. Our findings indicate that the gorillas effectively adapted to the changes, most likely by relying on social support, to navigate new conditions. This study contributes to our understanding of how socio-cognitively complex species cope with necessary alterations in captive animal care programs. Furthermore, these observations may inform strategies to enhance the welfare of zoo-housed animals and to improve their captive care.

## 1. INTRODUCTION

Promoting optimal captive animal welfare and quality of lives is of highest importance for institutions that are part of The World Association of Zoos and Aquariums (WAZA) and the European Association of Zoos and Aquariums (EAZA) (Kagan and Veasey, 2010). With novel research findings, evidence-based best practices and state-of-the-art technologies, the bar of acceptable standards for animal keeping is continuously raising higher (Maple, 2007), in terms of enclosure design and size, suitable behavioural enrichment, moving, socializing and training opportunities for animals, and adequate staff number (Ward et al., 2018). Changes in husbandry are often implemented to meet both animal welfare and operational needs. These changes may be physical, such as modifying the physical environment or relocating animals, or social, involving the introduction, removal or regrouping of individuals (Powell, 2010). Although such interventions are typically intended to benefit individuals in the long-term (Rose and Riley, 2022), they might represent short-term challenges in animal welfare (Morgan and Tromborg, 2007; Nevalainen, 2014; Teixeira et al., 2007). Case studies documenting relocations of particular captive individuals and groups provide valuable insights for caretakers, zoo managers and conservationists helping to minimize stress and negative emotional states such as fear, boredom or insecurity, while promoting positive welfare outcomes (Dawkins, 1990; Teixeira et al., 2007). These reports also enhance our understanding of how socio-cognitively complex animals cope with environment and social changes in captivity (Clark, 2017).

In non-human primates, relocating and changing group composition can lead to physiological changes (Cinque et al., 2017; Nehete et al., 2021; Schapiro et al., 2012), behavioral changes (Crockett et al., 2000, 1995) as well as changes in rates of social interactions (Ryan and Hauber, 2016). Transfers of great apes can be challenging particularly when moving to a new unfamiliar environment or separating from their previous social group. Indeed, reduced levels of activity and higher rates of scratching (i.e. potential behavioural indicator of stress) were reported in both gorillas and chimpanzees following relocation to a novel environment (Ross et al., 2011). However, such changes can also have positive effects by introducing new stimuli, social opportunities and enrichment, which can stimulate natural behaviours. For instance, in gorilla groups that regularly alternated between enclosures, increased activity levels were observed in new environment (Lukas et al., 2003) and they also decreased levels of undesirable behaviours (Goerke et al., 1987). Changes in social group composition can bring varied outcomes. Introduction of two bonobo groups increased affiliative interactions (Caselli et al., 2023), while spatial associations shifted after the integration of hand-reared gorilla infants (McCann and Rothman, 1999). In contrast, death of silverback did not result in significant behavioural changes in the group (Gartland et al., 2018). Temporarily elevated stress hormone levels have been reported both in individuals introduced to new group members (Jacobs et al., 2014) and in those remaining after others were relocated (Peel et al., 2005).

This study based on the Prague Zoo management decision to split the group of gorillas in two separate groups and relocate one of the subgroups to a new enclosure. The main study aim was to assess how these changes influenced gorilla behaviour. Based on the previous reports, we predicted gorillas would either i) reduce feeding, spend more time resting and less time moving as a consequence of potentially stressful changes represented by the split and relocation (e.g. Ross et al., 2011), or ii) increase locomotion, vigilance and exploration and decrease resting when dealing with the new conditions (e.g. Lukas et al., 2003). We also predicted that iii) potential indicators of stress and compromised welfare such as self-directed and ‘undesirable’ behaviours (i.e., coprophagy, hair plucking) will increase following the changes (Ross et al., 2011), whereas enrichment-related behaviour would decrease, with both returning to original levels when the animals habituate to new conditions.

## 2. METHODS

### 2.1. Subjects and housing

This study was conducted in Prague Zoo (Prague, Czech Republic). The initial group consisted of seven gorillas (see Table 1 for details): one adult silverback male, three adult females, two subadult and one juvenile males (classification based on Breuer et al. 2009). The group was housed in an enclosure (inside area 238 m2, outside area 811 m2) and a subgroup (see below 2.2.) relocated to a new enclosure (inside area 466 m2, outside area 2570 m2). Both enclosures were equipped with abundant climbing and resting structures and enrichment objects. The gorillas were fed a mixture of vegetables twice a day (at 8:30 and 14:30), with leafy and cruciferous vegetable spread across the enclosure twice a day (morning and afternoon) and with food pellets for gorillas (Marion Natural Foods) once a day. Water was available *ad libitum*.

**Table 1:**
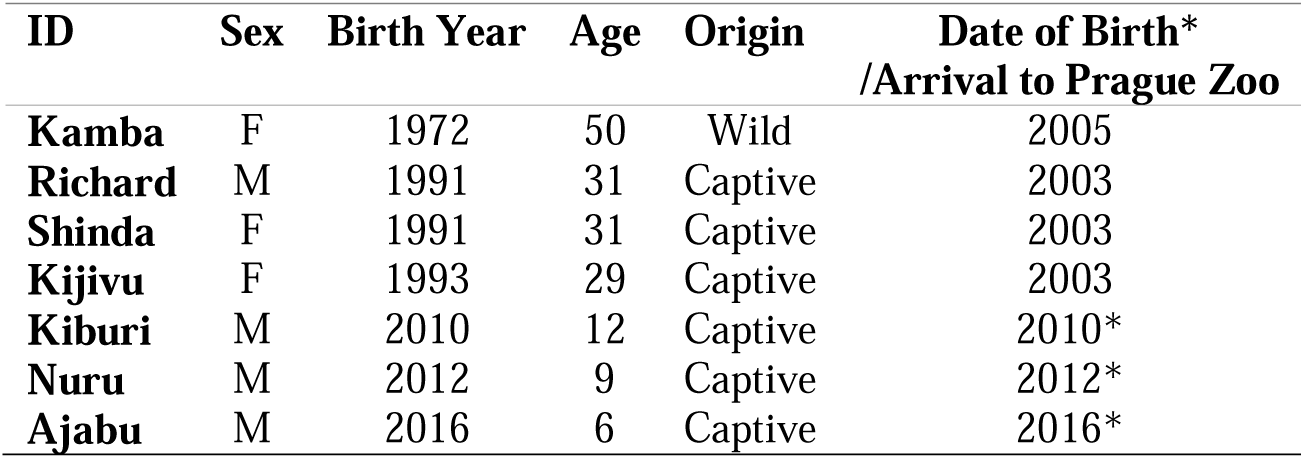
Subject information, with sex, year of birth, age at the time of the study, origin and date of birth or arrival to the Prague Zoo.

| ID | Sex | Birth Year | Age | Origin | Date of Birth*<br>/Arrival to Prague Zoo |
| --- | --- | --- | --- | --- | --- |
| **Kamba** | F | 1972 | 50 | Wild | 2005 |
| **Richard** | M | 1991 | 31 | Captive | 2003 |
| **Shinda** | F | 1991 | 31 | Captive | 2003 |
| **Kijivu** | F | 1993 | 29 | Captive | 2003 |
| **Kiburi** | M | 2010 | 12 | Captive | 2010* |
| **Nuru** | M | 2012 | 9 | Captive | 2012* |
| **Ajabu** | M | 2016 | 6 | Captive | 2016* |

### 2.2. Group split and relocation

The group split and relocation to a new enclosure were planned for two reasons; first, because the new enclosure provides more space and can accommodate a larger group of individuals, and second, because the zoo management needed to limit the reproduction of silverback male, as his offspring are overrepresented according to the international gorilla studbook. Thus, the zoo management decided to create two new groups: a group consisting of silverback male and two of his sons (Richard, Kiburi and Nuru, i.e. a ‘bachelor’ group) that would stay in the original enclosure and a group consisting of three females and the youngest juvenile male (Kamba, Shinda, Kijivu and Ajabu, i.e. a mixed-sex group) that would move into the new enclosure on June 8^th^, 2022. Following the changes, visitors were not allowed in the new enclosure during periods “after1” and “after2” (see 2.3. below) but were allowed in the old enclosure three weeks after the changes onwards. During the period “after1”, black and white colobus monkeys (*Colobus guereza*) join gorillas in the new enclosure. As the reaction of gorillas to this addition was negligible (main caretaker Martin Vojacek, personal communication) we did not split the data further to investigate the effect of this change. The detailed timeline of the observations and implemented changes are provided in Fig.1.

**Figure 1.**
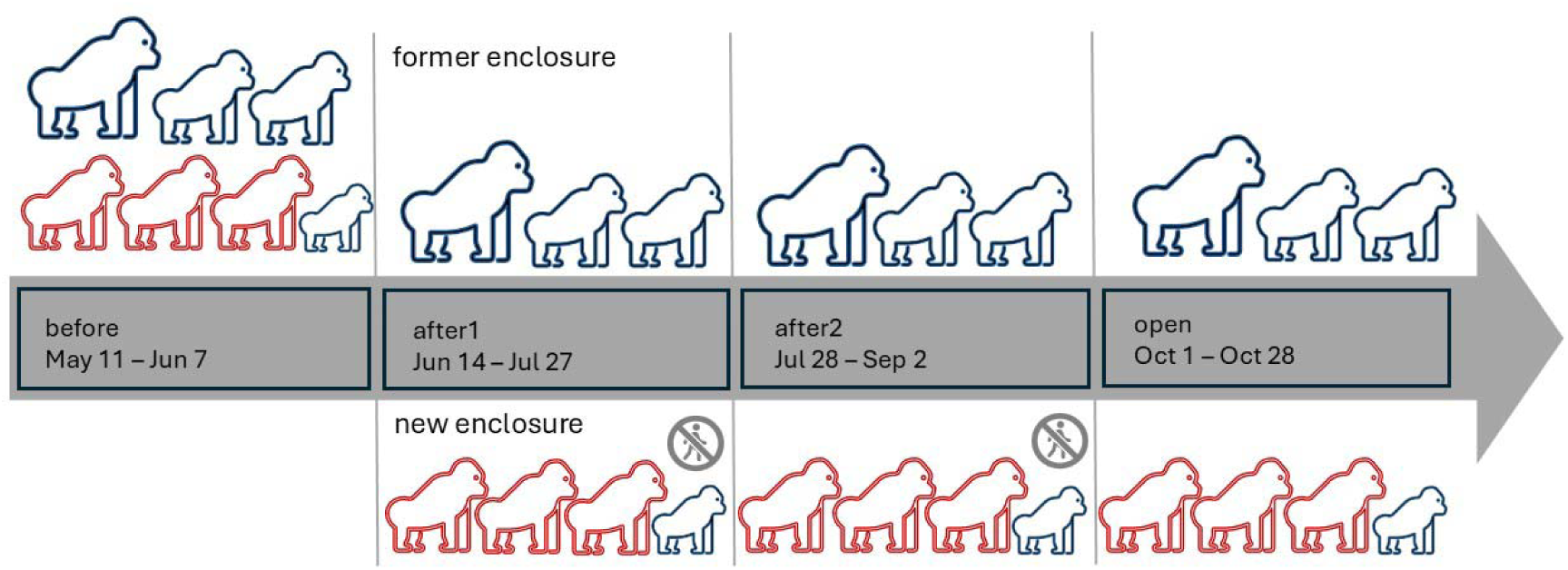
Timeline of study periods with main implemented changes: first observations occurred on May 11, group split and relocation on Jun 8, first observation after the changes in new enclosure on June 14, first observation after the changes in old enclosure on June 28, black and white colobus monkeys (*Colobus guereza*) were introduced to the new enclosure on July 14, new enclosure opened to public on September 28, and last observations were made on October 28.

In preparation for the move, the gorillas were occasionally trained: since April 2016 they were trained to enter the transport box, six months before the move, they were trained to the presence of the veterinarian and the silverback male was additionally trained to receive an injection in case drug administration was needed for the transport. At the time of relocation, three individuals (i.e. Richard, Nuru and Kijivu) were administered a sedative to avoid their anticipated negative reaction to the interventions. For one month after the group split, Richard was further treated with small doses of sedative to prevent severe socio-negative reactions (main caretaker Martin Vojacek, personal communication).

### 2.3. Data collection

Behavioral observations were done from May to October 2022, from 10 am to 6 pm each day, and were divided in four time periods: “before” (i.e. before the move; May 11- June 7), “after1” (i.e. immediately after the move; June 14 - July 27), “after2” (i.e. July 28 - September 2), and “open” (i.e. after the new enclosure was opened to public; October 1- October 28). The observations were split into “after1” and “after2” period to monitor the development of behavioural changes in time and to make periods comparable in terms of time and observations. The observations were conducted using a combination of focal instantaneous sampling and focal continuous sampling (Altmann, 1974) for 20-minute focal period with 2-minute sample points. Originally, we focused on 21 behavioural variables that covered daily activities, social interactions, self-directed and ‘undesirable’ behaviours (i.e., coprophagy, hair plucking, regurgitation), but 9 behaviours occurred rarely. Thus, only 12 variables were used in statistical analysis: resting, locomotion, feeding, social contact and proximity, approaches, grooming, scratch, groom-self, yawn, enrichment, and coprophagy. Behavioural variables recorded via instantaneous scan sampling were expressed as the proportion of the behavior in all scans, those recorded during continuous sampling were converted to frequencies per hour of observation, and those including social interactions (contact, proximity, approaches, grooming) were additionally divided by the number of social partners available, so they are expressed as number of interactions per social partner. This was done because there were fewer individuals to interact with in each of the two groups after the split than before the split. For the complete list of behavioral variables and their definitions see Table S1. Every individual was observed maximally twice per day (morning and afternoon). The observations were distributed evenly throughout the day and time periods, counterbalanced for different individuals, and were conducted from the visitor area. Two observers (MZ, MC) recorded data on separate days. Inter-observer reliability (IOR) was conducted on a set of two 30-minute videos using intra-class correlation (ICC) (Shrout and Fleiss, 1979). IOR was deemed sufficient if the ICC had values > 0.5, and p < 0.05. The agreement between two observers was excellent in the behaviours recorded by continuous (ICC (3, 1) = 0.917, [95% CI] = 0.721, 0.977, F = 21.1, p < 0.001) and instantaneous sampling (ICC (3, 1) = 0.967, [95% CI] = 0.889, 0.991, F = 62.5, p < 0.001).

Locomotion, feeding and resting behaviors were analysed to see if group split and move affected the daily time budget. Social interactions were analysed using proximity and contact, grooming and approaches. For these, we recorded the identity of the social partner and the directionality of the behavior. We also evaluated self-directed behaviors (i.e., scratch, self-groom, yawn), in enrichment involvement (i.e., attending to, manipulating and exploring any type of enrichment provided by the keepers) and stereotypical ‘undesirable’ behaviors as potential negative welfare/stress indicators (see Table S1 for details).

### 2.4. Statistical analysis

All statistical analyses were run in R v 4.0.5 (R Core Team, 2022). Although it is clear that the two new subgroups have different experiences as e.g. one was relocated and without visitors for some periods, we had to analyse them together as one group as separate analysis was not possible due to limited sample size (N=3 and N=4). The effect of time period on the behavioral variables (see below) was tested with general linear mixed models (GLMM) using the ‘glmmTMB’ package (Brooks et al., 2017). Response behavioural variables recorded via instantaneous scan sampling were analyzed using beta family and logit link. Behavioral variables recorded during continuous sampling were analyzed using Gamma family with log link if the data did not contain zeroes, or zero-inflated Gamma family if the data contained zeroes. In total, we ran 11 models with the following response variables: rest, feed, locomotion, proximity, approach, groom, scratch, self-groom, yawn, coprophagy, enrichment (details available in Table S1). The individual identity was set as a random effect and the period (“before”, “after1”, “after2”, “opening”) as a fixed effect. Significance of the fixed effect predictor was tested using the likelihood ratio tests using the *drop1* function. A post-hoc Tukey’s range test was performed if the results were significant. The model diagnostics were checked using the DHARMa package (Hartig, 2022) with diagnostic tests (Kolmogorov-Smirnov, dispersion, outlier) not significant (P > 0.05) in all models (details not shown).

## 3. RESULTS

### 3.1. Changes in locomotion, resting and feeding

In total 868 focal observations, i.e. 289.3 hours were collected with mean 10.8 hours (SD = 2.4) per individual per period. The locomotion time was significantly different among the studied periods (χ2 = 14.31, DF = 3, p = 0.003) (Fig. 2A): individuals spent significantly less time moving in the period before the move (i.e. “before”) in comparison to all other periods, although the comparison with “after1” period showed only a non-significant trend in this direction (Table 2). Time period had no significant effect on the proportion of time individuals spent resting (χ2 = 3.20, DF = 3, p = 0.362) (Fig. 2B), however time spent feeding differed significantly among the periods (χ2 =22.45, DF = 3, p < 0.001) (Fig. 2C). In particular, gorillas spent more time feeding in the period “open” in comparison to the first two periods (Tab. 3). Temporal trajectories of changes in locomotion, feeding and resting for each individual are depicted in Fig.3 A, 3B, 3C.

**Figure 2.**
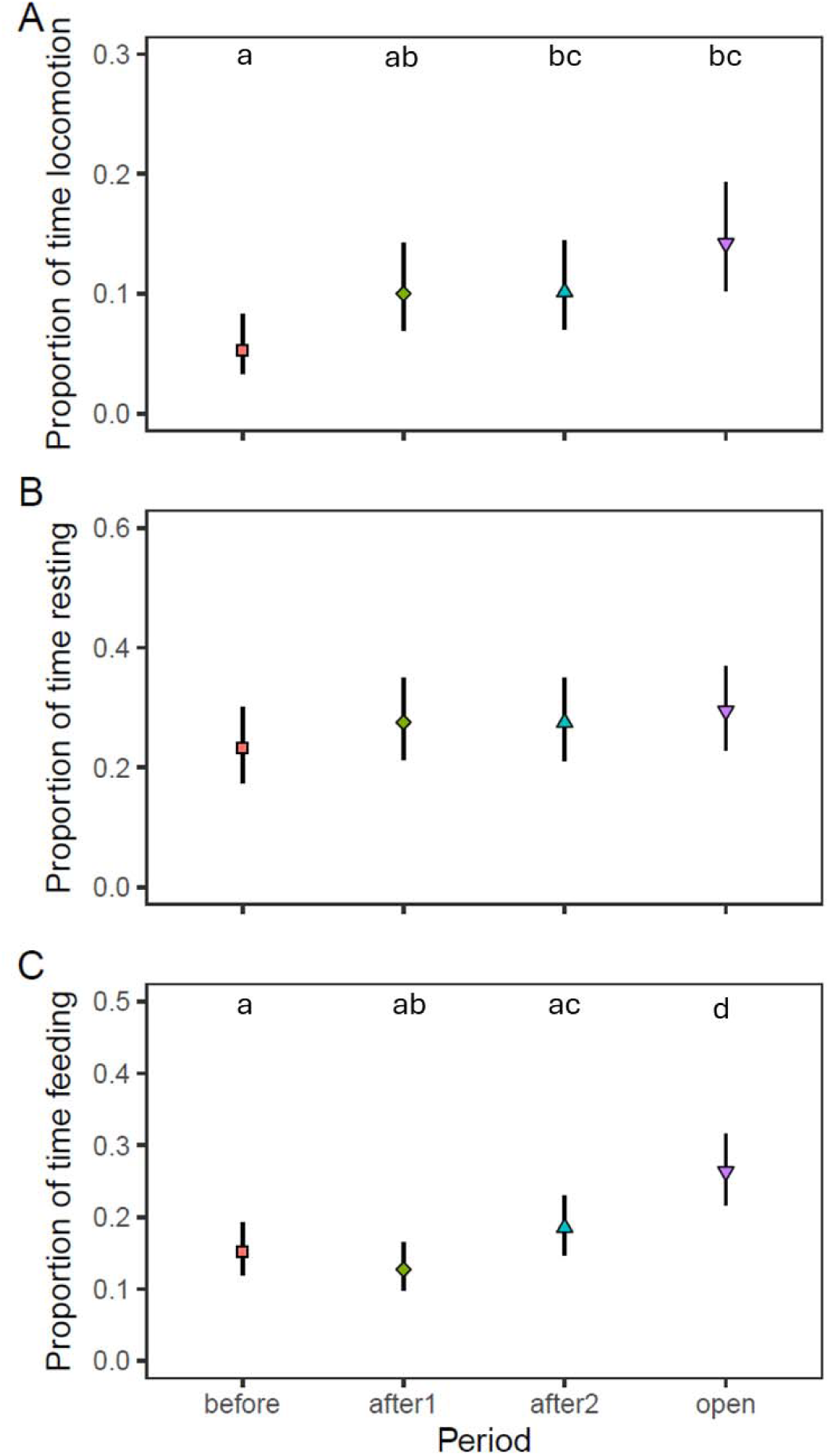
Comparison among the time periods in (A) proportion of time individuals spent moving, (B) resting and (C) feeding. Symbols = estimate from GLMM; error bars = 95% confidence intervals. Different letters indicate significant differences between periods (p<0.05).

**Figure 3.**
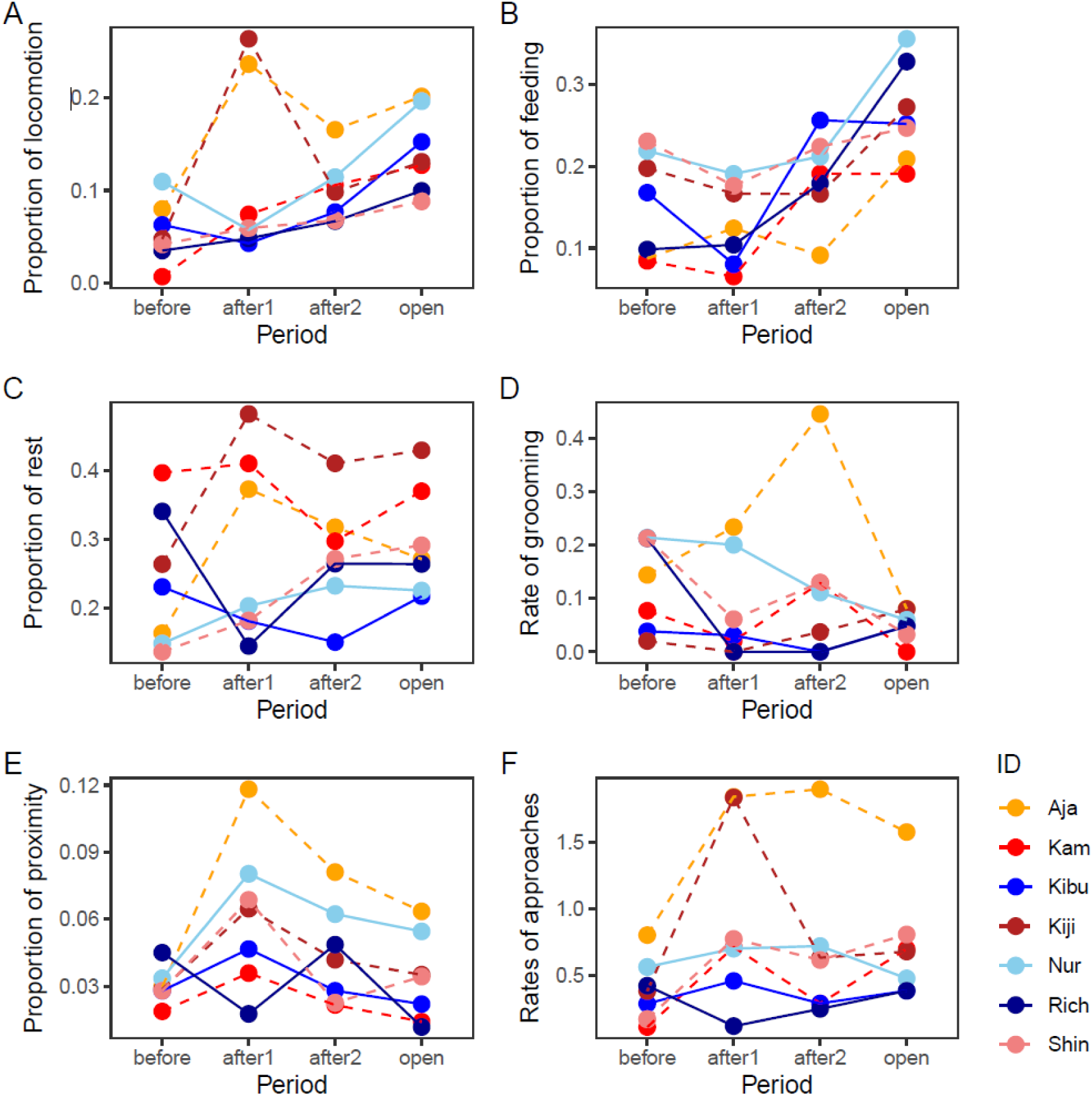
Temporal trajectories of individual proportion of time spent locomotion (A), feeding (B), rest (C), rates of grooming (D) proportion of time spent in proximity (E) and rates of approaches (F). Data shown as average proportion of instantaneous samples for each period (A, B, C, E) and as average rate per hour of observation for each period (D, F). Individuals from original enclosure are depicted in shades of blue with solid lines, individuals from new enclosure in shades of red with dashed lines.

**Table 2.** Results of the post-hoc Tukey’s range tests analyzing the differences between time periods for locomotion and feeding. P < 0.05 in bold.

| Response | Periods | Estimate | SE | z | P |
| --- | --- | --- | --- | --- | --- |
| Locomotion | after1 - before | 0.696 | 0.276 | 2.525 | 0.056 |
|  | **after2 - before** | 0.711 | 0.272 | 2.613 | **0.044** |
|  | **open - before** | 1.093 | 0.261 | 4.197 | **<0.001** |
|  | after2 - after1 | 0.014 | 0.236 | 0.061 | 0.999 |
|  | open - after1 | 0.397 | 0.223 | 1.777 | 0.282 |
|  | open - after2 | 0.382 | 0.220 | 1.738 | 0.302 |
| Feeding | after1 - before | -0.208 | 0.166 | -1.249 | 0.594 |
|  | after2 - before | 0.239 | 0.155 | 1.539 | 0.413 |
|  | **open - before** | 0.690 | 0.147 | 4.690 | **< 0.001** |
|  | **after2 - after1** | 0.446 | 0.162 | 2.759 | **0.029** |
|  | **open - after1** | 0.898 | 0.154 | 5.835 | **< 0.001** |
|  | **open - after2** | 0.452 | 0.142 | 3.192 | **0.008** |

**Table 3.** Results of the post-hoc Tukey’s range tests analyzing the differences between time periods for rate of approaches per hour and proportion of time spent in proximity. P < 0.05 in bold.

| Response | Periods | Estimate | SE | z | P |
| --- | --- | --- | --- | --- | --- |
| Approaches | **after1 - before** | 0.755 | 0.244 | 3.091 | **0.011** |
|  | after2 - before | 0.399 | 0.240 | 1.662 | 0.344 |
|  | open - before | 0.577 | 0.241 | 2.397 | 0.078 |
|  | after2 - after1 | -0.356 | 0.238 | -1.497 | 0.439 |
|  | open - after1 | -0.179 | 0.238 | -0.750 | 0.877 |
|  | open - after2 | 0.178 | 0.238 | 0.748 | 0.877 |
| Proximity | **after1 - before** | 0.745 | 0.184 | 4.058 | **< 0.001** |
|  | after2 - before | 0.388 | 0.191 | 2.031 | 0.175 |
|  | open - before | 0.136 | 0.202 | 0.673 | 0.907 |
|  | after2 - after1 | -0.357 | 0.163 | -2.190 | 0.125 |
|  | **open - after1** | -0.609 | 0.172 | -3.553 | **0.002** |
|  | open - after2 | -0.253 | 0.184 | -1.376 | 0.513 |

### 3.2. Changes in social interactions and social proximity

Grooming rate did not significantly change among the study periods (χ2 =5.68, DF = 3, p = 0.128) (Fig. 4A), however rate of approaches changed among the periods (χ2 =8.52, DF = 3, p = 0.036). Individuals approached each other more often in the period “after1” in comparison to the period “before” (Tab. 3, Fig. 4B). Time spent in proximity and in contact changed significantly among the study periods (χ2 =13.53, DF = 3, p = 0.004). Individuals spent more time in proximity to others during the first period after the move (i.e., “after1”) in comparison to periods “before” and “open” (Tab 3, Fig 4C). Temporal trajectories of changes in grooming, proximity and approaches for each individual are depicted in Fig. 3D, 3E and 3F.

**Figure 4.**
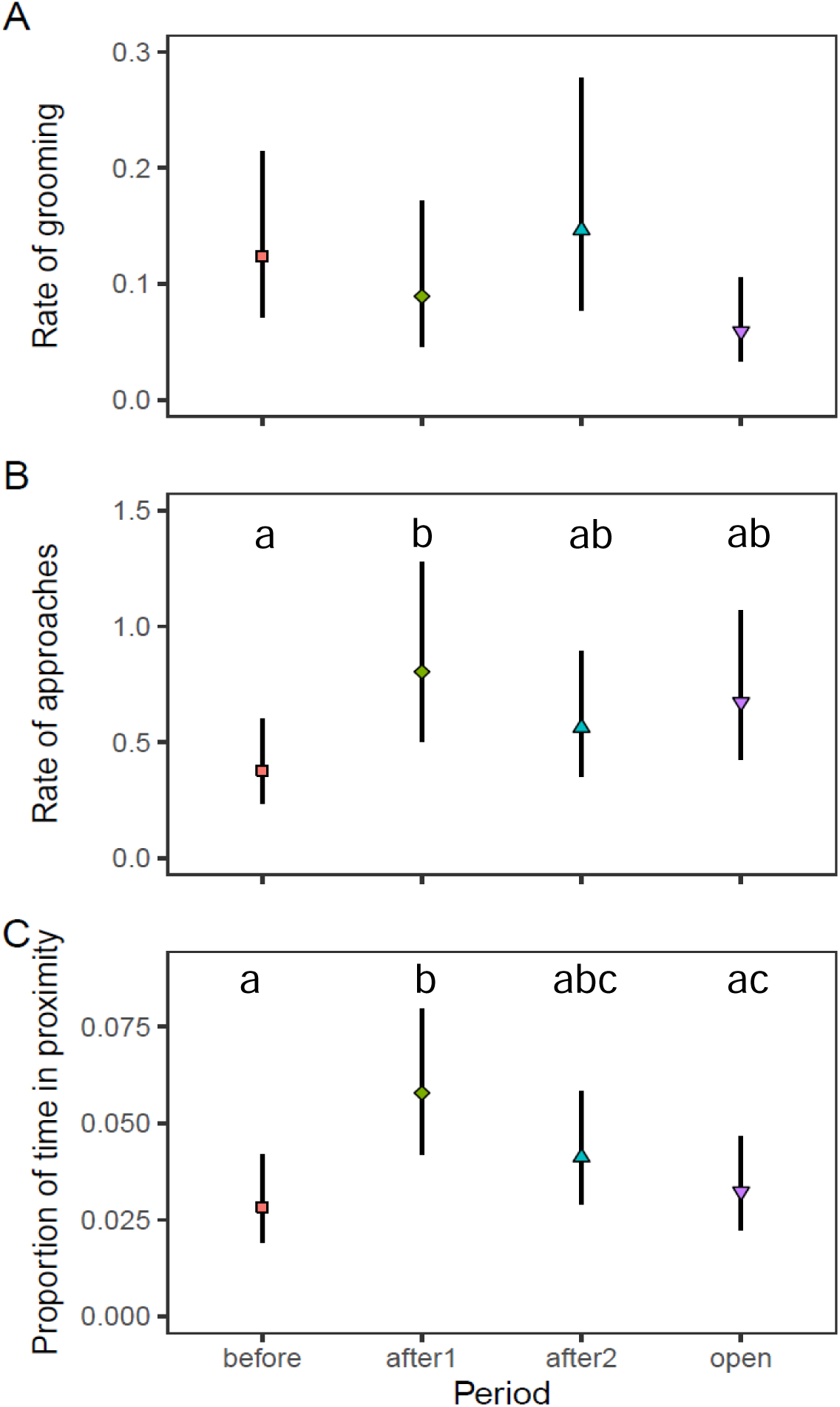
Comparison among the time periods in (A) rate of grooming initiated and received per hour per group member, (B) rate of approaches initiated and received per hour per group member and (C) proportion of time individuals spent in proximity or body contact per group member. Symbols = estimate from GLMM; error bars = 95% confidence intervals. Different letters indicate significant differences between periods (p<0.05).

### 3.3. Changes in self-directed, ‘undesirable’ and enrichment-related behaviours

Rate of scratching changed significantly among periods (χ2 =23.90, DF = 3, p <0.001): it decreased in the period “after1” in comparison to “before” and then further decreased throughout the study (Fig. 5A, Tab. 4). Rate of self-grooming showed a similar significantly decreasing pattern (χ2 =15.07, DF = 3, p =0.002) (Fig. 5B). There was no significant difference among time periods in the rates of yawning (χ2 =7.80, DF = 3, p =0.050) (Fig. 5C) and rates of enrichment-related behaviour (χ2 =6.91, DF = 3, p=0.075) (Fig. 5E), although there was a non-significant trend in both cases, caused by a single individual. In case of yawning, this effect was caused by Richard, the silverback male, whose rates of yawning sharply decreased after the group split. In case of enrichment-related behaviour, this effect was caused by Ajabu, the youngest juvenile male, whose rates of enrichment-related behaviour decreased after the group split and relocation. In other individuals there were no notable changes in either yawning nor enrichment involvement. Temporal trajectories of changes in scratching, self- grooming, yawning and enrichment-related behaviour for each individual are depicted in Fig. 6A, 6B, 6C and 6E.

**Figure 5.**
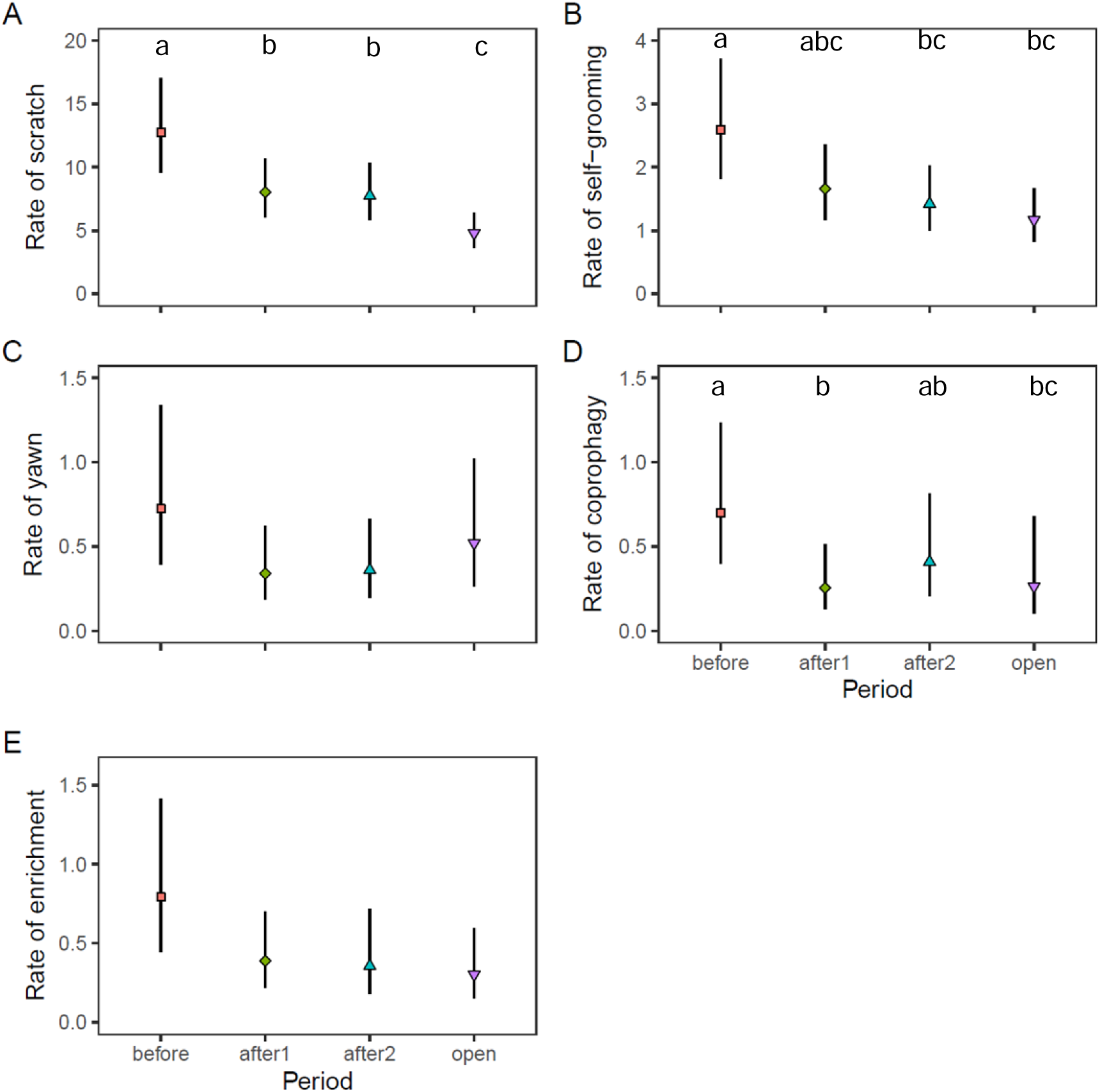
Comparison among the study periods in rates of (A) scratching, (B) self-grooming, (C) yawning, (D) coprophagy and (E) enrichment activity per hour of observation. Symbols = estimate from GLMM; error bars = 95% confidence intervals. Different letters indicate significant differences between periods (p<0.05).

**Figure 6.**
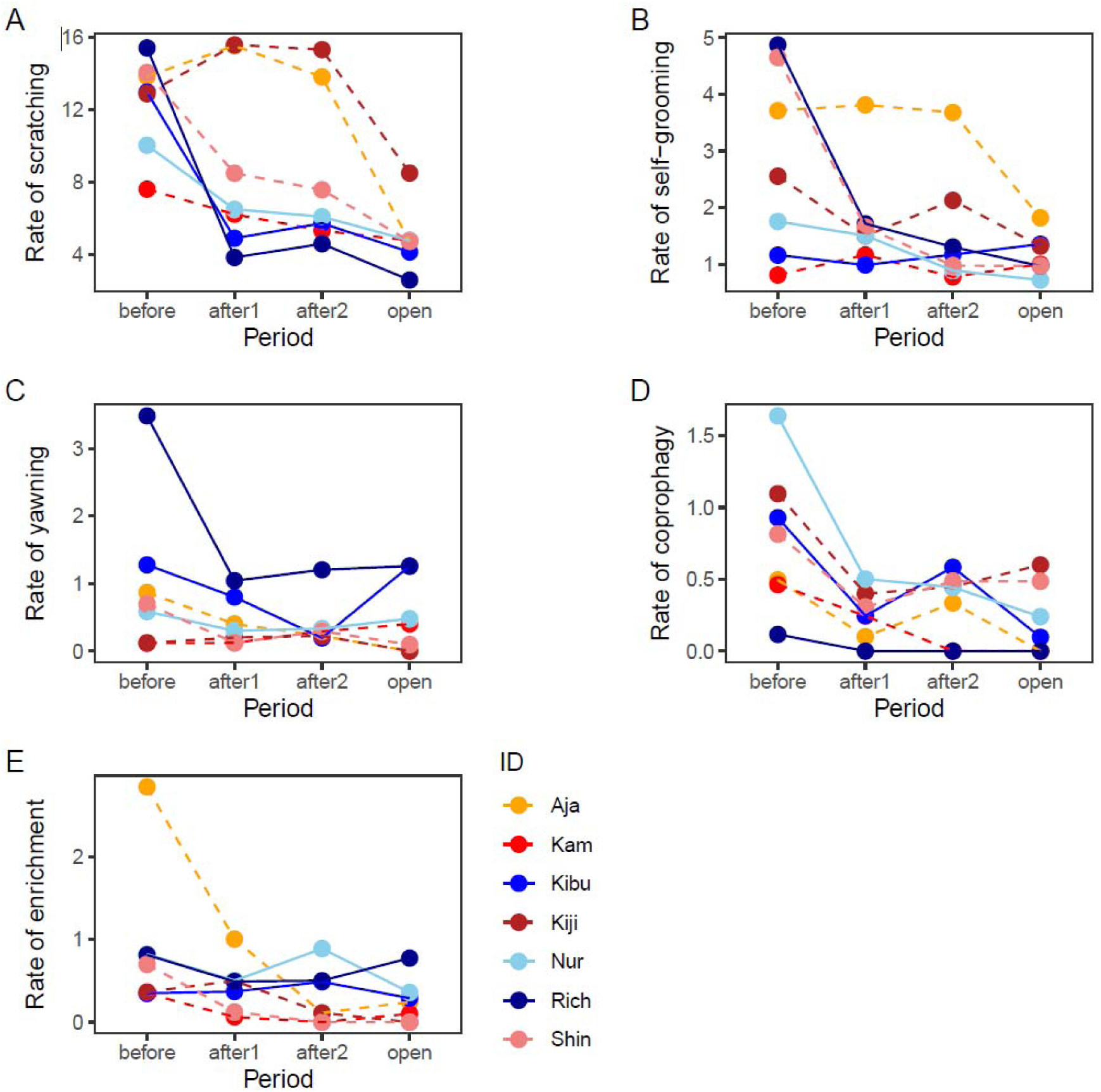
Temporal trajectories of individual rates of scratching (A), self-grooming (B), yawning (C), coprophagy (D) and enrichment involvement (E). Data shown as average rate per hour of observation for each period. Individuals from original enclosure are depicted in shades of blue with solid lines, individuals from new enclosure in shades of red with dashed lines.

**Table 4.** Results of the post-hoc Tukey’s range tests analyzing the differences between time periods for rates of scratching and self-grooming per hour. P<0.05 in bold.

| Response | Periods | Estimate | SE | z | P |
| --- | --- | --- | --- | --- | --- |
| Scratch | **after1 - before** | -0.463 | 0.146 | -3.175 | **0.008** |
|  | **after2 - before** | -0.498 | 0.145 | -3.434 | **0.003** |
|  | **open - before** | -0.978 | 0.145 | -6.766 | **< 0.001** |
|  | after2 - after1 | -0.035 | 0.143 | -0.248 | 0.995 |
|  | **open - after1** | -0.515 | 0.144 | -3.580 | **0.002** |
|  | **open - after2** | -0.480 | 0.144 | -3.339 | **0.005** |
| Self-groom | after1 - before | -0.448 | 0.186 | -2.414 | 0.074 |
|  | **after2 - before** | -0.603 | 0.187 | -3.229 | **0.007** |
|  | **open - before** | -0.801 | 0.188 | -4.258 | **< 0.001** |
|  | after2 - after1 | -0.155 | 0.184 | -0.842 | 0.834 |
|  | open - after1 | -0.353 | 0.185 | -1.910 | 0.224 |
|  | open - after2 | -0.198 | 0.185 | -1.069 | 0.708 |
| Coprophagy | **after1 - before** | -1.011 | 0.261 | -3.880 | **<0.001** |
|  | after2 - before | -0.540 | 0.268 | -2.012 | 0.180 |
|  | **open - before** | -0.979 | 0.362 | -2.708 | **0.034** |
|  | after2 - after1 | 0.471 | 0.274 | 1.720 | 0.308 |
|  | open-after1 | 0.032 | 0.322 | 0.100 | 0.999 |
| | open-after2 | -0.439 | 0.34\$ | -1.277 | 0.572 |

Some ‘undesirable’ behaviours (i.e., like coprophagy) significantly decreased after the group changes (χ2=10.49, DF=3, p=0.015) in all individuals (Fig. 5D, 6D). Other stereotypical ‘undesirable’ behaviors were rare in our dataset, which prohibited further statistical analyses. However, it is interesting to note that regurgitation, observed in three individuals before the move (with rates of 3.02, 0.62 and 0.24 per hour in period “before”), decreased substantially after the move-related changes (with rates of 0, 0.3, 0 per hour in period “after1”) and it was not recorded in any individual in the last period (i.e. “open”). One adult female often covered her ears in stressful situations, in rates of 4.9 per hour in the period “before”. This behavior decreased to the rates of 2.2, 0.3 and 0 per hour in the “after1”, “after2” and “open” periods, respectively. Contrary to this observation, hair plucking behavior in another adult female did not change much during the time periods (period “before” had 0.47 per hour, while period “open” had 0.29 per hour).

## 4. DISCUSSION

Our study on western lowland gorillas at Prague Zoo revealed significant alterations in their daily behaviours following a group split and relocation. After the changes, gorillas spent more time moving, approached each other more often and spent more time in proximity. A decrease in self-directed and ‘undesirable’ behaviours (i.e. behavioral indicators of negative welfare/stress) seem to indicate that the changes did not compromise gorilla welfare.

Daily activity of gorillas changed as a consequence of the environmental and social changes in both novel and original enclosures. Observed increased locomotion can be a sign of higher vigilance in novel environment but also indicates positive welfare (Lukas et al., 2003). The new enclosure was larger and more complex, while the original one housed fewer individuals after the changes and both of these factors likely promoted more opportunities for physical activities. Resting patterns remained unchanged, but surprisingly feeding time increased notably in the last phase (“open”). This increase is difficult to interpret as the change in feeding was most evident in the final period of the study not around the implemented changes. Though, seasonal changes in energy intake and activity levels were documented in both wild and captive gorillas (Masi et al., 2015; Ross et al., 2011). Given the several months of duration of our study the seasonal changes in feeding could have influenced our findings.

Both the time spent in proximity and the approach rates increased in the period after the changes (“after1”) and slowly decreased in the later stages of the study (“after2”, “open”) returning slowly to pre-change levels. This pattern can be seen as a coping strategy of gorillas in the new potentially challenging situation, where they might seek company of others for comfort or reassurance. Changes in social networks and increased time spent in proximity have been documented in previous studies of gorillas following social changes (Hoff et al., 1996; McCann and Rothman, 1999), often driven by infants and juveniles (Hoff et al., 1996). Indeed, in our study, the juvenile male Ajabu showed most pronounced increases of social proximity and approaches. These findings are in agreement with social buffering phenomenon (Kikusui et al., 2006), where the mere presence of conspecifics mitigates stress (Denommé and Mason, 2022; Kikusui et al., 2006). Thus, the changes in the social spacing reflect the challenging nature of the situation for the gorillas but also represent one of possible mechanism how to cope with them. In contrast, grooming rates remained unchanged, corroborating findings of Hoff and colleagues (1998) when the natural social change represented by the death of silverback did not cause measurable changes in grooming behaviour (Hoff et al 1998), and despite their otherwise known role in stress reduction and social bonding (Boccia et al., 1989; Shutt et al., 2007). No changes of average grooming rates in our study may be explained by contradictory individual tendencies (Fig XX). In particular, the adult silverback male Richard experienced a drop in grooming rates due to relocation of females, his main grooming partners, whereas the youngest juvenile male Ajabu experienced an increase in grooming rates, probably seeking social comfort from others (i.e. due to having more difficulties to adjust to the new conditions; main caretaker Martin Vojacek, personal communication). These individuallly specific responses highlight the value of detailed behavioural monitoring that might allow to provide specific welfare support to particular individuals based on their predispositions (Gartland et al., 2018).

Although modern animal keeping facilities are striving to reduce various potential sources of stress (Morgan and Tromborg, 2007), some inevitable changes may trigger a stress response, like elevated cortisol levels in relocations of great apes (Behringer et al., 2012; Schapiro et al., 2012). In our study, behavioural markers of stress and negative welfare decreased and remained low during the study suggesting a positive adaptation to the changes. Enclosure rotation (Lukas et al., 2003) or subject relocation to more natural enclosures lead to decrease of self-directed behaviors (a traditional proxy for stress and negative welfare assessment) in gorillas and chimpanzees (Clarke et al., 1982; Ross et al., 2011). For instance, regurgitation fully disappeared in several individuals and Kijivu’s ear-covering behavior observed usually under stresfull conditions (Peel et al., 2005), notably decreased. The new social conditions for the bachelor group, especially the lack of females, might have reduced stress-related behaviour, particularly that of the silverback male. This male held a tenure position for twenty years, more than three times longer than the wild average (Breuer et al., 2010). Increased stress levels due to changes can also be quite brief and can return to baseline within several days (Cinque et al., 2017; Peel et al., 2005). In our study, we were not allowed to observe the gorillas for a few days immediately after the group split and relocation, as the zoo prioritized calm conditions for the gorillas. If a brief stress spike occurred, it may have gone undocumented.

Careful planning to allow sufficient training time can also help to minimize negative effects of changes, as training is also seen as a form of enrichment and gives the animals more sense of control and predictability (Westlund, 2014). Long-term and targeted pre-interventions training likely helped to reduce the stress levels in gorillas. Furthermore, continued housing with known social partners in both subgroups and remaining within the same facility with known caretakers may have contributed to no observable increase in stress-related behaviour after the changes (Schaffner and Smith, 2005).

While it is overall good to reduce stress, changes and challenging situations are also a natural part of animals’ lives in the wild. Both sexes disperse from their natal group and join other groups and thus adapt to new social and environmental conditions (Forcina et al., 2019). They regularly navigate the shifting group dynamics and unfamiliar areas (Breuer et al., 2010; Stokes, 2004). When managed carefully, similar changes in captivity can be enriching and more reflective of natural experiences than living in conditions of limited novelty and variation.

This case study has several limitations, mostly due to management decisions. The study was initially designed to monitor the relocation of the entire group, but the zoo management subsequently decided to also split the group, introducing two major changes, i.e. move to a new enclosure and a group split. This makes the interpretation of the behavioural changes difficult in relation to the importance of the two changes. Due to the low sample size, meaningful statistical comparisons, between the relocated mixed-sex group and non-relocated bachelor group was not feasible. However, visual inspection of the individual temporal trajectories of behavioral changes, reveal no notable group differences (Figs. XX-XX), with an exception that the bachelor group showed a more pronounced increase in feeding and a steeper sustained decline in scratching. As the two groups also differed in sex composition, it remains unclear whether the observed differences were sex-related or due to relocation.

Although scratching and self-directed behaviours have been widely used as stress and negative-welfare indicators in various species (Cairo-Evans et al., 2022), including gorillas (Carder and Semple, 2008; Ross et al., 2010), some studies have not found a consistent link (Ross et al., 2010). This suggest that current markers may not fully capture responses to complex challenges. Further studies could explore alternative metrics such as behavioural diversity or enclosure use variability (Fernandez et al., 2023).

Visitors’ presence also varied: the original enclosure was mostly accessible throughout the study, while the new enclosure remained closed to visitors during the “after1” and “after2” periods. While some studies report number of visitors influenced ape behaviour (Birke, 2002; Carder and Semple, 2008; Cook and Hosey, 1995), others do not (Hosey et al., 2023).

The group of seven gorillas at Prague Zoo is slightly above the zoo average (usually six) (ZIMS Species Holdings, 2024), it is still a small sample. Larger sample size would enable more targeted statistical analysis, but would likely require long-term, multi-zoo collaboration effort (Fowler et al., 2022; Garcia-Pelegrin et al., 2022).

Despite these limitations, this case study offers valuable insights into how captive gorillas respond to environmental changes, including group split and relocation. By carefully monitoring the effects of such changes on behaviour of captive individuals and individual variations in their coping mechanisms, researchers and zoos can better understand and support the welfare of captive great apes.

## Ethical statement

The study was non-invasive and purely observational and adhered to legal requirements of Czech Republic. Approval to conduct the study was granted by the Prague Zoo. No approval was required from the authors’ institutions.

## CRediT authorship contribution statement

Marie Žabková: Investigation, Data Curation, Writing - Original Draft: Writing - Review & Editing Monika Čížková: Investigation, Writing - Review & Editing, Vedrana Šlipogor: Methodology, Writing – Original Draft, Writing - Review & Editing, Roman Vodička: Conceptualization, Resources, Martina Konečná: Conceptualization, Methodology, Formal analysis, Writing - Original Draft, Writing - Review & Editing, Supervision

## Acknowledgements

We thank the staff of the Prague Zoo and specifically to Martin Vojáček and all gorilla keepers. We are especially grateful to director of Prague Zoo Miroslav Bobek, who authorized our study, and to František Vácha, who initiated the cooperation on this project. Thanks to David Boukal for statistical advice, help with figure preparations and comments on the manuscript.

