## Supplementary material for "Smooth move: Behavioural changes in captive western lowland gorillas (*Gorilla gorilla gorilla*) after group split and relocation"

Table S1: Ethogram of recorded behaviours. Instantaneously recorded behaviours were noted down every two minutes. Continuously recorded behaviours were recorded any time they occurred during the 20minute focal period.

| Name | Behavioral category | Continuous/  Instantaneous | Analysed as | Definition |
| --- | --- | --- | --- | --- |
| feeding | general | I | proportion of scans | An animal is consuming food (and/or enrichment, branches, leaves), including chewing and food manipulation |
| resting | general | I | proportion of scans | An animal is sitting or laying, eyes are closed for most of the time, no obvious activity |
| grooming | social | C/I | rate per hour of observation, adjusted for number of social partners available | An animal is going through hair of the other, while watching the groomed place on the other‘s body, using its fingers or mouth, may or may not pick up some particles |
| grooming-self | self-directed | C/I | rate per hour of observation | An animal is going through hair of its own body, is watching the groomed place, may or may not pick up some particles |
| locomotion | general | I | proportion of scans | Any kind of movement resulting in changing its position and which is not defined as part of another behaviour, including walking/running/climbing/jumping for a distance longer than 3 meters, or which results in changing the substrate (e.g. coming from tree log to the ground) |
| contact | social | I | proportion of scans, adjusted for number of social partners available | An animal is in direct physical contact with another conspecific, i.e. touching with any part of the body, but not grooming with the conspecific, or doing any other defined behaviour |
| proximity | social | I | proportion of scans, adjusted for number of social partners available | An animal is out of direct contact but within an arm’s reach of another conspecific |
| coprophagy | undesirable | C/I | rates per hour of observation | An animal feeds on faeces (its own or of another conspecific) |
| enrichment | welfare indicator | C/I | rates per hour of observation | An animal attends to any type of enrichment provided by the keepers, includes object manipulation, play and exploration |
| scratching | self-directed | C | rates per hour of observation | Animal is going fast through its fur and body surface with its arm/hand, leg/foot, or is rubbing self against another surface, no visible attention to the scratched part |
| yawning | self-directed | C | rates per hour of observation | An animal clearly opens its mouth in automatic manner |
| approach | social | C | rates per hour of observation, adjusted for number of social partners available | An animal comes into proximity (an arm’s reach) of the other or others |
| departure | social | C | rates per hour of observation, adjusted for number of social partners available | An animal goes out of proximity (an arm’s reach) of the other or others |
| hair-plucking | undesirable | C | rates per hour of observation | A single (or multiple) hair is plucked with a rapid jerking-away motion, may be accompanied by inspection and consumption of the hair shaft and follicle, can be self-directed or done to another individual |
| regurgitation | undesirable | C | rates per hour of observation | An animal voluntarily brings up partially digested food and then reingests it; considered as abnormal behaviour |
| vigilance | general | I | not analysed | Animal is stationary in any posture while paying attention (i.e. monitoring) its environment or actively scanning the environment, can also watch a particular stimulus |
| social play | welfare indicator | C/I | not analysed | Physical play with a partner, includes chase, mock bite, poke/hit, throw at, object tug, wrestle, etc, and excluding any aggressive events. Does not necessitate continuously maintained proximity with a play partner |
| solitary play | welfare indicator | C/I | not analysed | Physical play without a partner, includes locomotory play and object play |
| contact aggression | social | C | not analysed | Charge another individual with bite, push, grab/pull hair, hit/slap, poke, wrestle |
| non-contact aggression | social | C | not analysed | Charge another individual with hitting or throwing object(s), chase, roar |
